# Economic costs of biological invasions in the United States

**DOI:** 10.1101/2021.07.02.450757

**Authors:** Jean E. Fantle-Lepczyk, Phillip J. Haubrock, Andrew M. Kramer, Ross N. Cuthbert, Anna J. Turbelin, Robert Crystal-Ornelas, Christophe Diagne, Franck Courchamp

## Abstract

The United States has thousands of invasive species, representing a sizable, but unknown burden to the national economy. Given the potential economic repercussions of invasive species, quantifying these costs is of paramount importance both for national economies and invasion management. Here, we used a novel global database of invasion costs (InvaCost) to quantify the overall costs of invasive species in the United States across spatiotemporal, taxonomic, and socioeconomic scales. From 1960 to 2020, reported invasion costs totaled $4.52 trillion (USD 2017). Considering only observed, highly reliable costs, this total cost reached $1.22 trillion with an average annual cost of $19.94 billion/year. These costs increased from $2.00 billion annually between 1960-1969 to $21.08 billion annually between 2010-2020. Most costs (73%) were related to resource damages and losses ($896.22 billion), as opposed to management expenditures ($46.54 billion). Moreover, the majority of costs were reported from invaders from terrestrial habitats ($643.51 billion, 53%) and agriculture was the most impacted sector ($509.55 billion). From a taxonomic perspective, mammals ($234.71 billion) and insects ($126.42 billion) were the taxonomic groups responsible for the greatest costs. Considering the apparent rising costs of invasions, coupled with increasing numbers of invasive species and the current lack of cost information for most known invaders, our findings provide critical information for policymakers and managers.

## Introduction

Biological invasions damage natural systems worldwide (Pyšek et al. 2020, Simberloff 2015). Non-native invasive species, organisms introduced beyond their natural range by human activity, can cause negative ecological and economic impacts as they spread through the novel environment. These species degrade ecosystem services (e.g., Walsh et al. 2016), disrupt natural communities (e.g., Dorcas et al. 2012), and significantly threaten or endanger native species (Blackburn et al. 2019). These damages are exemplified by a number of species that individually have had massive, widely apparent impacts. In the United States, well-publicized examples include zebra and quagga mussels (*Dreissena polymorpha* and *D. bugensis*, respectively), which have altered biophysical characteristics in the Great Lakes and clogged water intakes (e.g., Miehls et al. 2009), Burmese pythons (*Python bivittatus*), brown tree snakes (*Boiga irregularis*), and rats (*Rattus* sp.) which have reduced or extirpated native birds, reptiles, and mammals (Dorcas et al. 2012, Wiles et al. 2003, Doherty et al. 2016), salt cedar (*Tamarix* spp.) which has disrupted surface and groundwater in the western US (Zavaleta 2000), the emerald ash borer (*Agrilus planipennis*) which has reshaped tree communities, particularly in urban areas (Kovacs et al. 2010), and the chytrid fungal disease (*Batrachochytrium dendrobatidis*) which has severely reduced many amphibian populations (Dueñas et al. 2018).

The wide-reaching ecological impacts of individual invasive species in the United States have been correlated with marked economic impacts. For instance, zebra mussels cost businesses and communities over $5 billion in the first 10 years after invasion alone (Boelman 1997) and emerald ash borers have been estimated to cost $10 billion over a decade in lost forest resources (Kovacs et al. 2010). Collectively, invaders threaten US agriculture (Paini et al. 2016), damage critical infrastructure (e.g., water treatment facilities, electrical power; Boelman 1997, Connelly et al. 2007), and substantially lower the value of property and other personal assets (Johnson and Meder 2013). For example, rush skeletonweed (*Chondrilla juncea*) caused an estimated $62 million annual loss to wheat, potato, legume, and hay crops in Washington state alone (Mefford et al. 2017). Further, rapidly spreading invasive insects, such as the gypsy moth (*Lymantria dispar*) and hemlock wooly adelgid (*Adelges tsugae*) threaten forest resources and human wellbeing (Mcmanus and Csóka 2007, Aukema et al. 2011), while expanding invasive mosquito populations vector pathogens that cause massive human health costs (Shepard 2011).

Given the great diversity in types of economic impacts from invasive taxa in the United States, the actual aggregate costs remain highly uncertain. This uncertainty is mainly due to the lack of synthesis of costs reported across spatiotemporal, taxonomic, and socioeconomic scales. This can result in highly variable cost estimates, sometimes differing by orders of magnitude. For instance, feral cats (*Felis catus*) cause great damage to native species, and multiple attempts have been made to value the effects of their depredations. However, these attempts vary widely, ranging from $30/bird (Pimentel et al. 2005) to $1500/bird (Lohr et al. 2013). When multiplied by the hundreds of millions of birds killed by cats every year, the discrepancy between estimates is vast. In an example of scale mismatch, Anderson et al. (Anderson et al. 2016) quantified agricultural damage costs of wild pigs (*Sus scrofa*) across 11 US states based on empirical observations, while Systma and Rouhe (2007) extrapolated control costs of pigs within a single state in a different decade. Overall, the disparate nature of cost estimates, combined with the lack of centralized systems that denote attributes such as method reliability, complicates cost comparisons across contexts. Ultimately, the thousands of invasive species recorded in the United States represent a sizable burden to the country’s economy, but the extent of this burden is unknown.

While economic impacts of a few individual invasive species have been estimated across the entire US (e.g., Martin and Blossey 2013), there are no current, comprehensive estimates of total costs. The most recent estimate of gross economic costs for the United States was $120 billion per year in 2005 (Pimentel et al. 2005), but this was criticized for methodological shortcomings, such as extrapolations from unclear baselines and a lack of spatiotemporal granularity (Hoffman and Broadhurst 2016, Cuthbert et al. 2020). Extrapolated and uncertain cost estimates are particularly problematic in the context of the US economy, given its size and importance within the global economy. Indeed, the United States has the largest economy in the world, with a 2019 GDP of $21 trillion (World Bank 2020) and is amongst the top three global importers/exporters, including trade agreements with 75 countries worldwide (Office of the United States Trade Representative 2020). With the increasingly open economy, global trade volume has risen markedly over the past 50 years. Costs due to invaders are thus likely increasing across the United States (as they are globally, Diagne et al. 2021), as higher trade volumes introduce a suite of new species, while climate change facilitates the establishment and spread of already introduced species (Seebens et al. 2017, 2018, Lockwood et al. 2019). Such increases, coupled with the current lack of reliable cost appraisals, inhibit effective decision making by policy makers involved in prevention and management of biological invasions in the United States, as well as hamper effective communication of the problem to the general public. While not all impacts of invasive species are economically quantifiable, robust estimates of their economic impacts can be a convincing way of communicating the scale of the problem to a diverse audience. Thus, a synthesized, comprehensive record of invasion costs is urgently needed to highlight the necessity of invasive species management to both decision makers and the public.

The InvaCost database (Diagne et al. 2020a) seeks to address this lack of robust cost information by presenting a comprehensive global database of reported costs of invasive species. It links costs from a variety of source documents with standardized taxonomic, sectorial, regional, and temporal descriptors. We used this database, as well as complementary cost sources, to synthesize and analyze currently available information on costs of invasions in the US economy. Specifically, our aim was to quantify how these costs are distributed by region, cost type, environment, societal sector, and taxonomic group, as well as to calculate annual and cumulative costs of invasive species from 1960 to present. Quantifying these values provides a vital step towards understanding the true socioeconomic impact of invasive species across the United States and implementing efficient and evidence-based management actions.

## Methods

### InvaCost database

To estimate the costs invasive species have on the US economy, we extracted recorded costs for the United States from the InvaCost database, which consists of 9,823 entries from 1605 studies of reported economic costs of invasive species (Diagne et al. 2020a, https://doi.org/10.6084/m9.figshare.12668570). The data included in the InvaCost database were retrieved via a structured review of publications found in the ISI Web of Science platform, Google Scholar, the Google search engine and through consultation with invasive species experts (Diagne et al. 2020a), along with analogous searches conducted in more than 10 non-English languages (e.g., French, Spanish, Chinese, and Japanese; Angulo et al. 2021; https://doi.org/10.6084/m9.figshare.12928136). Individual cost records were converted to USD 2017. We examined the resulting database subset for double-counted and redundant data. All such redundancies were removed from the database or edited to remove overlapping dating of costs, as appropriate. All corrected costs were forwarded to as requested by InvaCost managers to update the main database. The resulting dataset contained 1534 entries specific to the United States, derived from 416 studies (Dataset S1).

To derive the costs of invasive species on the US economy, we filtered the database to include country “USA”, thereby excluding costs shared between the United States and other countries (e.g., Canada). We added an additional descriptor to the database that classified the entries by region: Northeast, Southeast, Midwest, Southwest, West, Outlying territories (Puerto Rico, Guam, Commonwealth of the Northern Mariana Islands, minor atolls), and multiregional/unspecified. We aimed to provide the most robust, yet conservative estimates of the costs of invasions in the US. For this purpose, we analyzed only highly reliable and observed costs, using the Invacost database method reliability and implementation type categorizations. Assessing methods for estimating costs across hundreds of heterogeneous studies reporting on diverse taxonomic groups, habitats, and economic sectors, makes a clear dichotomy between high and low reliability very challenging (Diagne et al. 2021). To maximize objectivity, the study used well defined, consistent criteria to assess study reliability (see Diagne et al. 2020a for full details on these criteria). Highly reliable entries were those classified in the ‘Method reliability’ column as *high*, and include data from peer-reviewed, official, and repeatable materials, as opposed to *low*, non-verifiable or non-repeatable estimates. Observed costs were those classified in the ‘Implementation’ column as an *observed*, actually realized cost, as opposed to a *potential* cost, which is not currently accrued or realized. As a result, we excluded from analysis all entries that were not from peer-reviewed literature or official reports (e.g., government documents), or were otherwise not reproducible, as well as those that were extrapolated but not empirically observed (Diagne et al. 2020a). The resulting estimate, derived from the best available data, represents a robust summation of reported costs of invasive species in the United States, which is, in turn, a minimum estimate of actual costs, as many costs likely go unestimated and unreported.

In addition to overall invasion costs, we also analyzed costs by several key components, both across the entire US and by region. These include the variables of cost type, environment, impacted sector, and taxa (see Table 1 for details and category definitions). Cost types were categorized into broad impact categories within the InvaCost database (Diagne et al. 2020a). For each variable, if the respective criteria were unspecified or covered multiple categories, we gave it an aggregate category (mixed cost type, mixed environment, multisectoral, multitaxon).

**Table 1.**
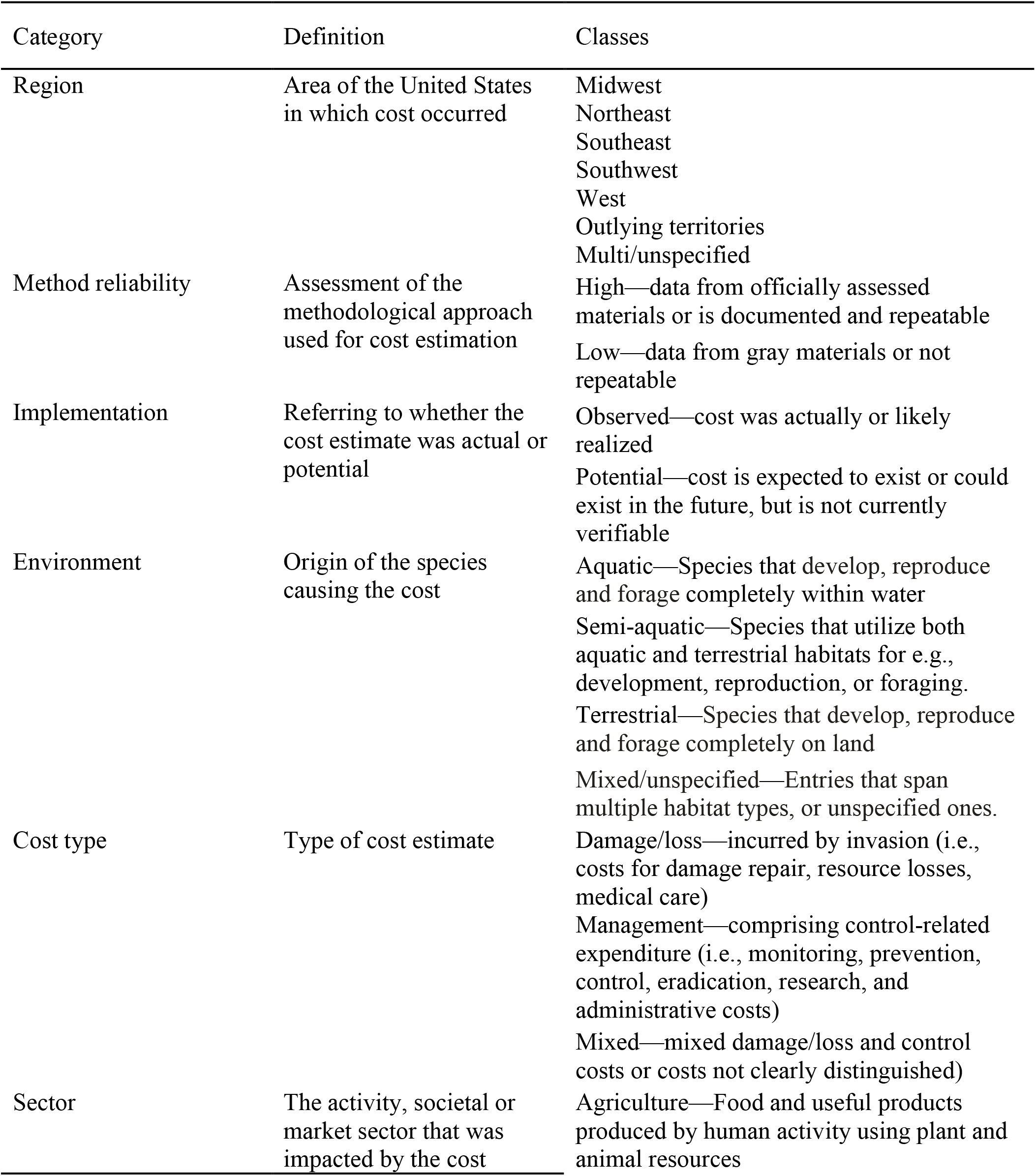
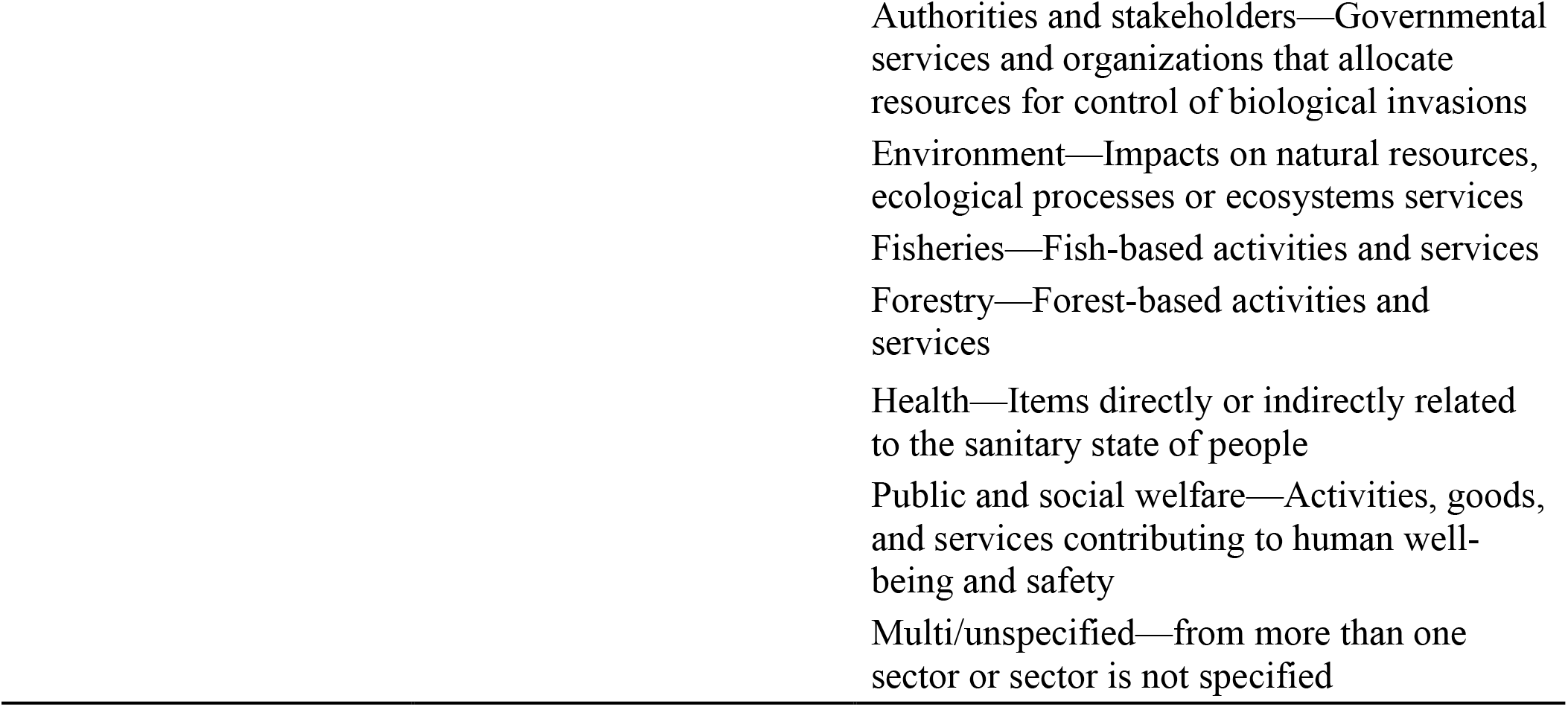
Categories, definitions and classes of variables included in the InvaCost database. Definitions of categories and classes are from Diagne et al. 2020a.

| Category | Definition | Classes |
| --- | --- | --- |
| Region | Area of the United States in which cost occurred | Midwest<br>Northeast<br>Southeast<br>Southwest<br>West<br>Outlying territories<br>Multi/unspecified |
| Method reliability | Assessment of the methodological approach used for cost estimation | High—data from officially assessed materials or is documented and repeatable<br>Low—data from gray materials or not repeatable |
| Implementation | Referring to whether the cost estimate was actual or potential | Observed—cost was actually or likely realized<br>Potential—cost is expected to exist or could exist in the future, but is not currently verifiable |
| Environment | Origin of the species causing the cost | Aquatic—Species that develop, reproduce and forage completely within water<br>Semi-aquatic—Species that utilize both aquatic and terrestrial habitats for e.g., development, reproduction, or foraging.<br>Terrestrial—Species that develop, reproduce and forage completely on land<br>Mixed/unspecified—Entries that span multiple habitat types, or unspecified ones. |
| Cost type | Type of cost estimate | Damage/loss—incurred by invasion (i.e., costs for damage repair, resource losses, medical care)<br>Management—comprising control-related expenditure (i.e., monitoring, prevention, control, eradication, research, and administrative costs)<br>Mixed—mixed damage/loss and control costs or costs not clearly distinguished) |
| Sector | The activity, societal or market sector that was impacted by the cost | Agriculture—Food and useful products produced by human activity using plant and animal resources |

### Costs over time

We analyzed average annual costs and cumulative costs using yearly cost data output from the *expandYearlyCosts* function in the ‘invacost’ package in R v4.0.2 (R Core Team 2020, Leroy et al. 2020). This function calculated annual costs (hereafter called annual cost entries) by dividing the total cost after conversion to USD 2017 by the number of years over which the total cost was incurred. We therefore calculated the cost over time for an individual entry by determining when the cost first and last occurred for every entry in our database, using information from the document reporting the cost. For example, a total cost of $100,000 reported by the original source as accruing over ten years would correspond to ten entries of $10,000 each. Specifically, we derived the total cumulative cost of invasions over time by calculating the probable duration time of each cost entry (duration time = probable end year of cost - probable first year of cost). When no starting year was indicated, we conservatively used the reference’s publication year. In some cases, probable ending year information was missing for potentially ongoing costs, which are costs likely to be repeated over years (contrary to one-time costs occurring only once). When no period of impact was specified, we counted only a single year (though costs might have repeated over many years, even to the present), making these estimates conservative, but also contributing to high variance between years.

We further used the costs over time to quantify the average annual costs of invasives in the United States between 1960 and 2020, and the average annual cost by decade over this period. Given the known time lags between the actual occurrence of costs and their reporting in the literature, we used quantiles from the time difference between when reported costs occurred and when they were published to apply time lag adjustments (25% = 1 year, 50% = 4 years, 75% = 11 years; Leroy et al. 2020, Diagne et al. 2021). We thus applied a correction to account for incomplete years, whereby, based on the above quantiles, costs after 2016 were removed from analysis as our model predicted < 50% of expected costs have been reported. We then fit the temporal dynamics of reported costs with generalized additive models (GAM), multivariate adaptive regression splines (MARS), robust regression to reduce the influence of outliers, least squares regression using the sandwich variance adjustment, and quantile regression (quantiles 0.1, 0.5, 0.9) all as implemented by default in the *costTrendOverTime* function from the ‘invacost’ package (Leroy et al. 2020). Model performance was assessed with root mean square error (RMSE; lower values indicate a better fit), with the understanding that not all models are equally appropriate for the data subset, which necessitates some qualitative rationale in model choice (Leroy et al. 2021). In evaluating models, we made the qualitative assumption that costs due to invaders are most likely stable or increasing because invasion rates worldwide show no sign of saturation (Seebens et al. 2017), and thus economic impacts from invasive species are unlikely to be decreasing. At the same time, both awareness and reporting of economic impacts of non-native invaders are rising (Diagne et al. 2021), making it even more likely that any recent declines in costs would be due to time lags in reporting, rather than actual decreases in costs.

## Results

Reported invasion costs across the US from 1960 to 2020 totaled $1.22 trillion (n = 1,750 annual cost entries) when conservatively considering only observed, highly reliable cost estimates. When we considered all data, reported costs reached $4.52 trillion (n = 4,790). Of these, 52% ($2.36 trillion; n = 2,645) were observed and 48% ($2.16 trillion; n = 2,145) were potential costs. Most costs (62%) originated from highly reliable sources ($2.78 trillion; n = 3,432), while $1.73 trillion (n = 1,357; 38%) were classified as low reliability. For the purposes of this study, we opted to be conservative and only focused on observed, highly reliable costs for further analysis.

Across regions within the United States, invasion costs differed considerably (Fig. 1; Table S1). The West had the highest regionally defined costs ($28.84 billion, n = 747) with over half of the estimated region-specific costs (58%), followed by the Southeast ($17.31 billion; n = 211), the Southwest ($2.06 billion; n = 26), the Midwest ($1.11 billion; n = 59), the outlying territories ($1.04; billion; n = 94), and lastly the Northeast ($632.30 million; n = 77). However, the vast majority of costs ($1.17 trillion; n = 536) were from multiple or unspecified regions.

**Fig. 1.**
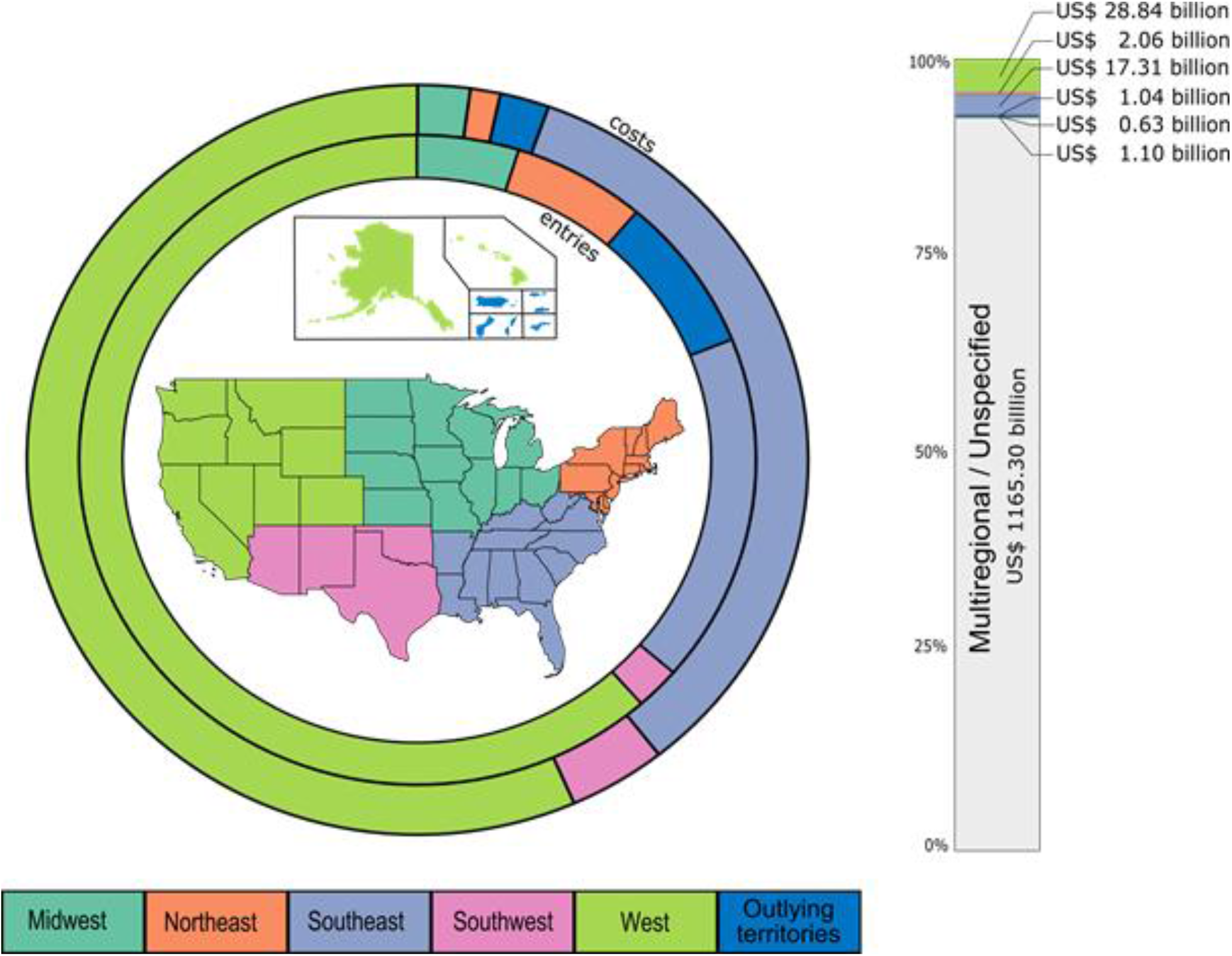
Shares of regionally defined costs and database entries for the six regions of the United States (1960-2020); costs and entries that were multiregional/unspecified were not included in the map as they could not be assigned to a specific region. Costs are represented by the outer circle and number of database entries by the inner circle. Taxon icons indicate the costliest (outer circle) and most reported species (inner circle) per region. The adjacent bar includes multiregional and unspecified region estimates.

### Cost types and environments

Over two-thirds (73%) of reported costs (see Table 1 for cost type definitions) across the United States were related to resource damages and losses ($896.22 billion; n = 647), while management costs were $46.54 billion (4%; n = 718). The remaining 23% ($273.52 million; n = 385) were mixed or unspecified costs (Fig. 2). Damage costs were also highest in most regions except for the Southwest, where mixed costs dominated (Fig. 2, Table S2). The fraction of reported costs arising from management was highly variable, ranging from as high as 33% in the outlying territories to 2% in the Southeast.

**Fig. 2.**
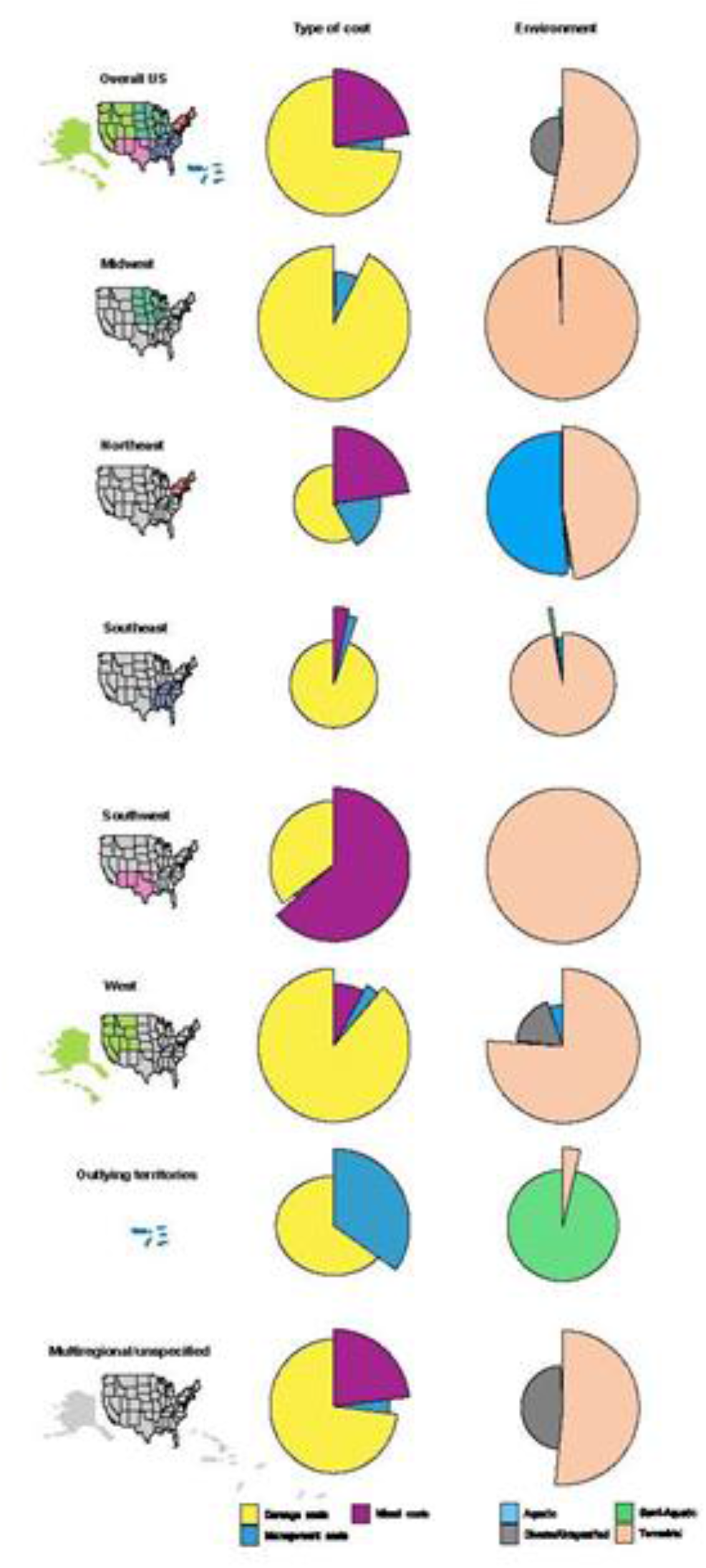
Distribution of cost types and impacted environments across US regions (1960-2020). The shares of each cost or environment type are scaled according to their respective number of entries in the database. Radius length represents the number of entries and arcs show the proportion of costs.

With regard to environment types (see Table 1 for definitions of environment types) across the entire United States (Fig. 2), invaders from terrestrial environments ($643.51 billion; n = 1,156; 53%) caused the majority of reported costs, followed by those from aquatic environments ($13.45 billion; n = 189; 1 %), and semi-aquatic environments ($1.57 billion; n = 193; < 1%). Entries from mixed or unspecified environments contributed $557.75 billion (n = 212; 46%). This held true within the US regions, where terrestrial costs dominated in all regions except the Northeast and the outlying territories, followed by costs in aquatic and semi-aquatic environments (Fig. 2, Table S3).

### Sectors and taxa

When activity sector (see Table 1 for sector definitions) was defined, agriculture was the most impacted ($509.55 billion, n = 259), followed by environmental ($102.59 billion; n = 108), forestry ($42.61 billion; n = 20), public and social welfare sectors ($40.74 billion, n = 166), and authorities-stakeholders ($37.11; n = 839). Fewer costs were reported for the health sector ($19.49 billion; n = 61), while fisheries ($40.12 million; n = 7) was the least impacted sector in our dataset. Distributions of costs within sectors differed considerably across regions. Of regionally specific costs, agriculture was the most impacted sector in the Midwest and West, public and social welfare in the Northeast and the Southwest, and forestry in the Southeast, while multisectoral costs dominated in the outlying territories (Fig. 3, Table S1).

**Fig. 3.**
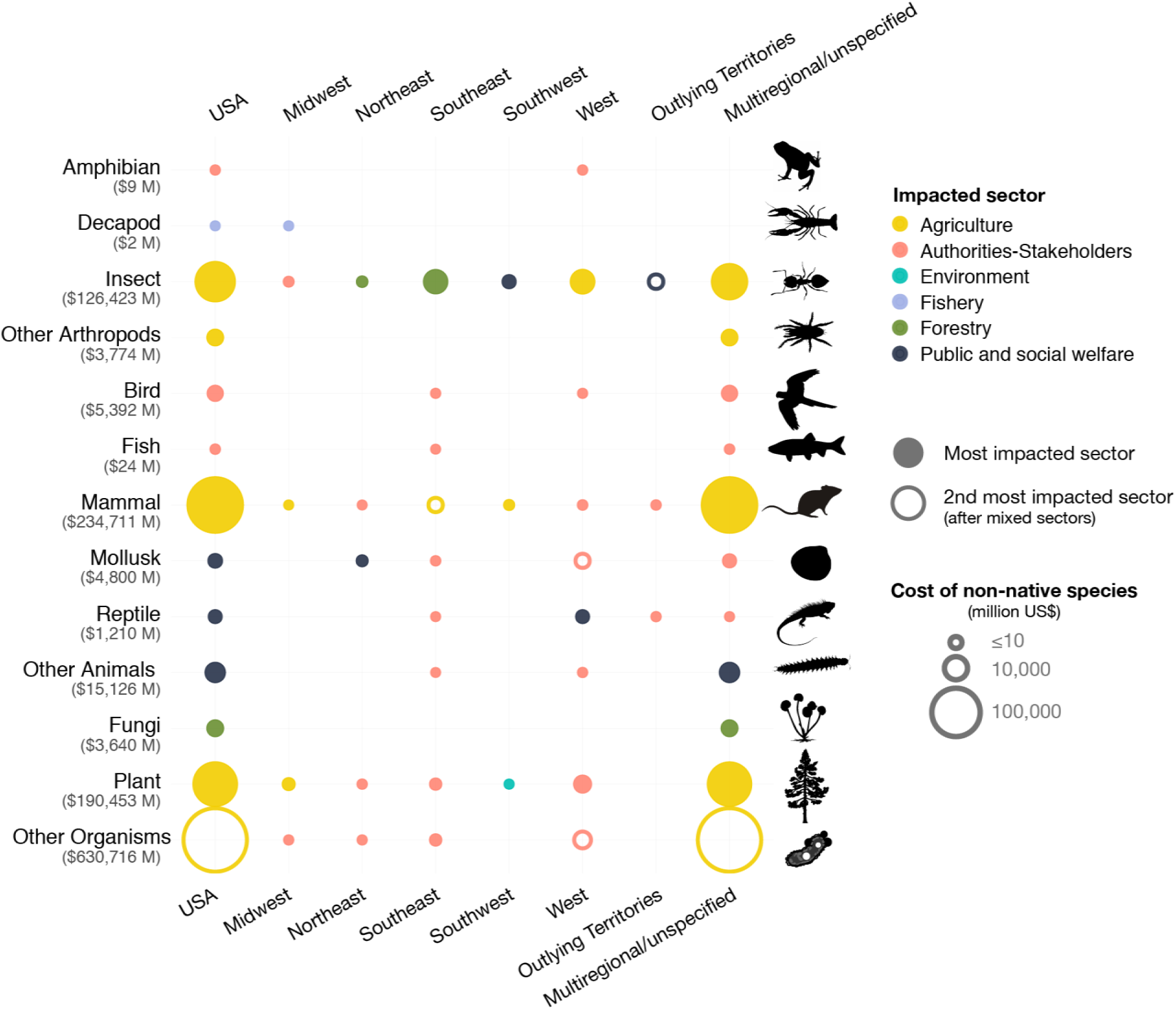
Magnitude of cost impacts (1960-2020) by broad taxonomic groups across regions. Filled color circles indicate which sector is most impacted by a specific group in a given region and unfilled color circles show the second most impacted sector when mixed sectors (costs attributed to more than one sector) are most impacted. For example, in the Southeast, insects were most costly to the forestry sector, while in the Southwest, insects costs were the highest for mixed sectors and second highest for agriculture. The “other organisms” category includes bacteria, chromista, viruses and costs associated with species from different kingdoms (e.g., plantae and animalia).

Across the United States as a whole, mammals were the costliest class of invaders ($234.71 billion) with the agriculture, environment, and mixed sectors primarily bearing the costs (Fig. 3). Plants were the second costliest invaders at $190.45 billion impacting primarily the agriculture sector. This was followed by insects($126.42 billion), birds ($5.39 billion), mollusks ($4.80 billion), fungi ($3.64 billion), reptiles ($1.21 billion), fish ($24.36 million) and amphibians ($9.29 million), with decapods reporting the lowest group-specific costs at $1.89 million. Other arthropods (excluding insects and decapods) contributed $3.77 billion, other animals contributed $15.13 billion, and other organisms, including microorganisms and undefined taxa $630.72 billion. Where regions were identified, insects and plants continue to be within the top three costliest groups in all regions, except for the outlying territories where only insects are amongst the top three (Fig. 3). Within the US regions, sectors impacted by invader classes also showed some key differences, with the Northeast showing primary impact by mollusks in the public and social welfare sector, the Southeast by insects in the forestry sector, the Midwest by plants in the agricultural sector, the West and Southwest by insects in the agricultural sector, and the outlying territories by insects in the public and social welfare sector (Fig. 3)

Six of the ten species with the highest reported, observed costs were insects, and nine of the ten species were animals, with Dutch elm fungus (*Ophiostoma ulmi*) the only exception (Table S4). Costs identifiable to specific species made up $326.49 billion, with most reported costs being contributed by multiple or unspecified species (Fig. 3). We also identified regional differences in cost contributing species, such that there was little overlap in species with the greatest reported costs from region to region (Table 2), though across regions, animals were the dominant species (12 vs. 3 species, respectively). Only the feral pig, brown tree snake, and red imported fire ant appeared in the top 3 most expensive species in more than one of the region-specific species lists.

**Table 2.**
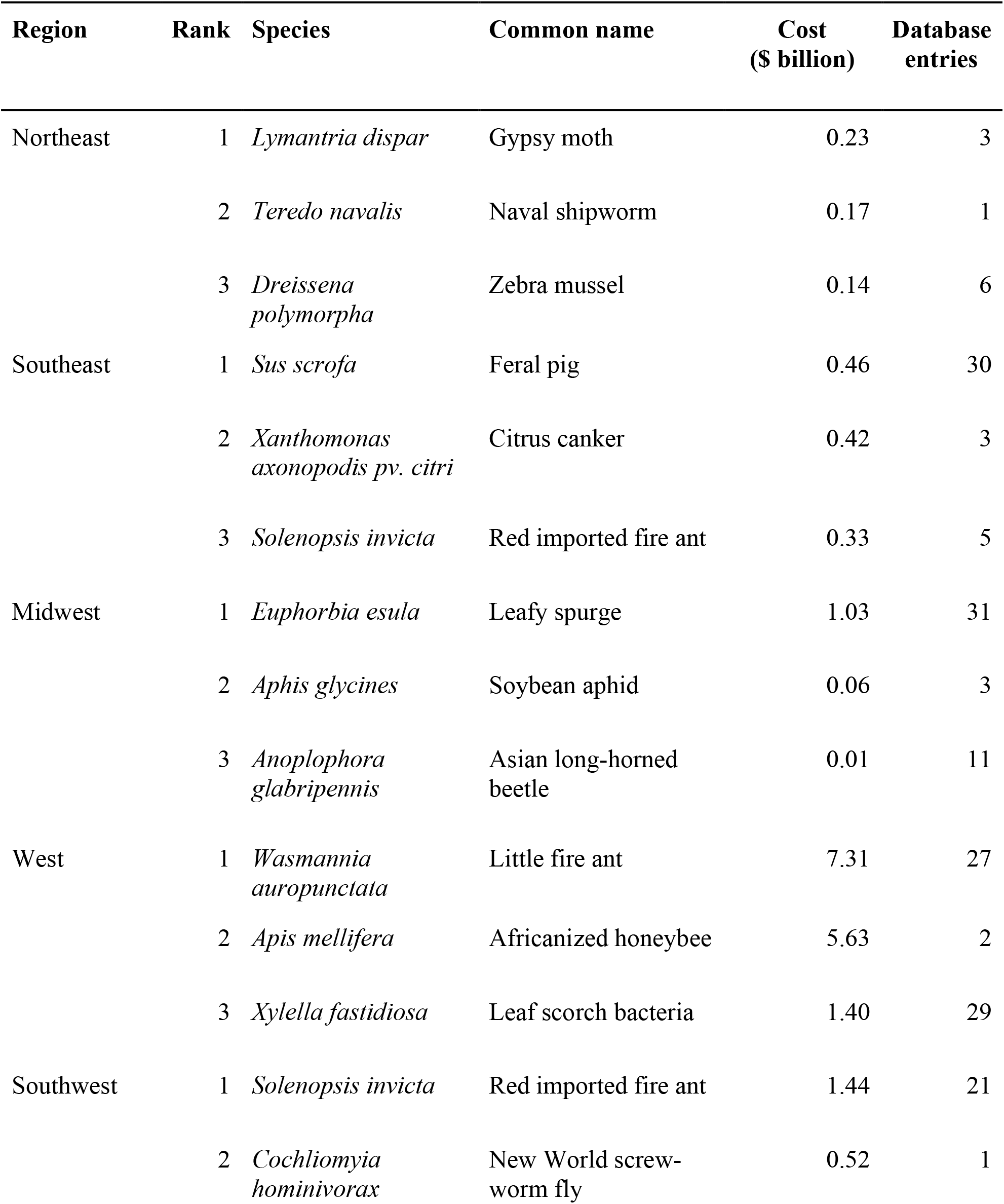
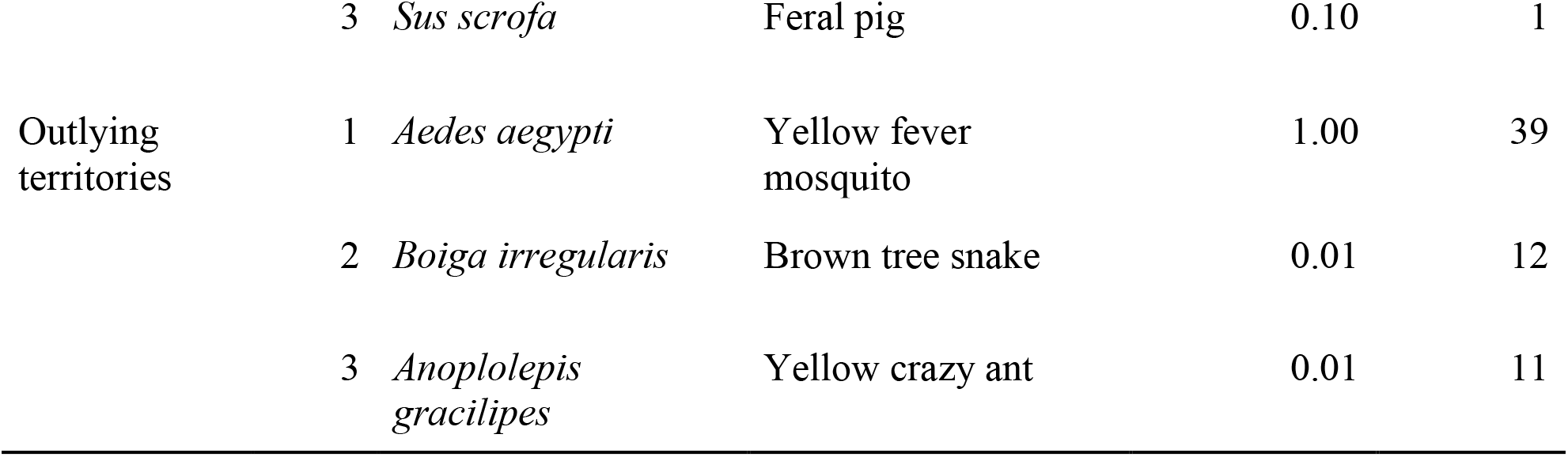
Top 3 species with highest reported costs per region in the United States. Numbers of associated database entries are included.

### Costs over time

The recorded total cost of $1.22 trillion between 1960 and 2020 (Fig. S1) amounted to an average annual cost over the entire period of $19.94 billion per year. This cost increased from $2.0 billion per year between 1960-1969 to $21.08 billion per year between 2010-2020. Models differed in their predictions of invasive species costs borne by the US economy over the 1960– 2020 period (Fig. 4; Table S5). The GAM predicted costs of $4.01 billion in 2020, while robust regression predicted 2020 costs at $6.19 billion (linear) and $0.28 billion (quadratic). The latter projected decreasing costs in recent years, which may be caused by a sensitivity to time lags in reporting, and it also had a relatively poor fit to the data (Table S5). The RMSE between the GAM and linear model was competitive (ΔRMSE = 0.02) (Table S5) and error bounds were large, reflecting high cost variance in recent years, which favored the use of robust regression over ordinary least squares regression. The estimate provided by quantile regression ranged over orders of magnitude, with the 0.1st quantile at $0.54 billion, the 0.5th quantile at $3.97 billion, and 0.9th quantile at $107.32 billion in 2020. The MARS model provided the lowest RMSE (Table S5), however it exhibited high sensitivity to recent results, creating an apparently spurious decrease in recent costs. In fact, although both the more flexible non-linear models (MARS and quadratic robust regression) had lower or comparable RMSE, this was driven by under-reported costs in recent years. For these models, predicted current costs fell below the overall annual average due to extremely high variance in recent costs and the occurrence of the three highest assigned costs in the 2000s, a probable artifact of the incompleteness of recent costs due to time lags in reporting.

**Fig. 4.**
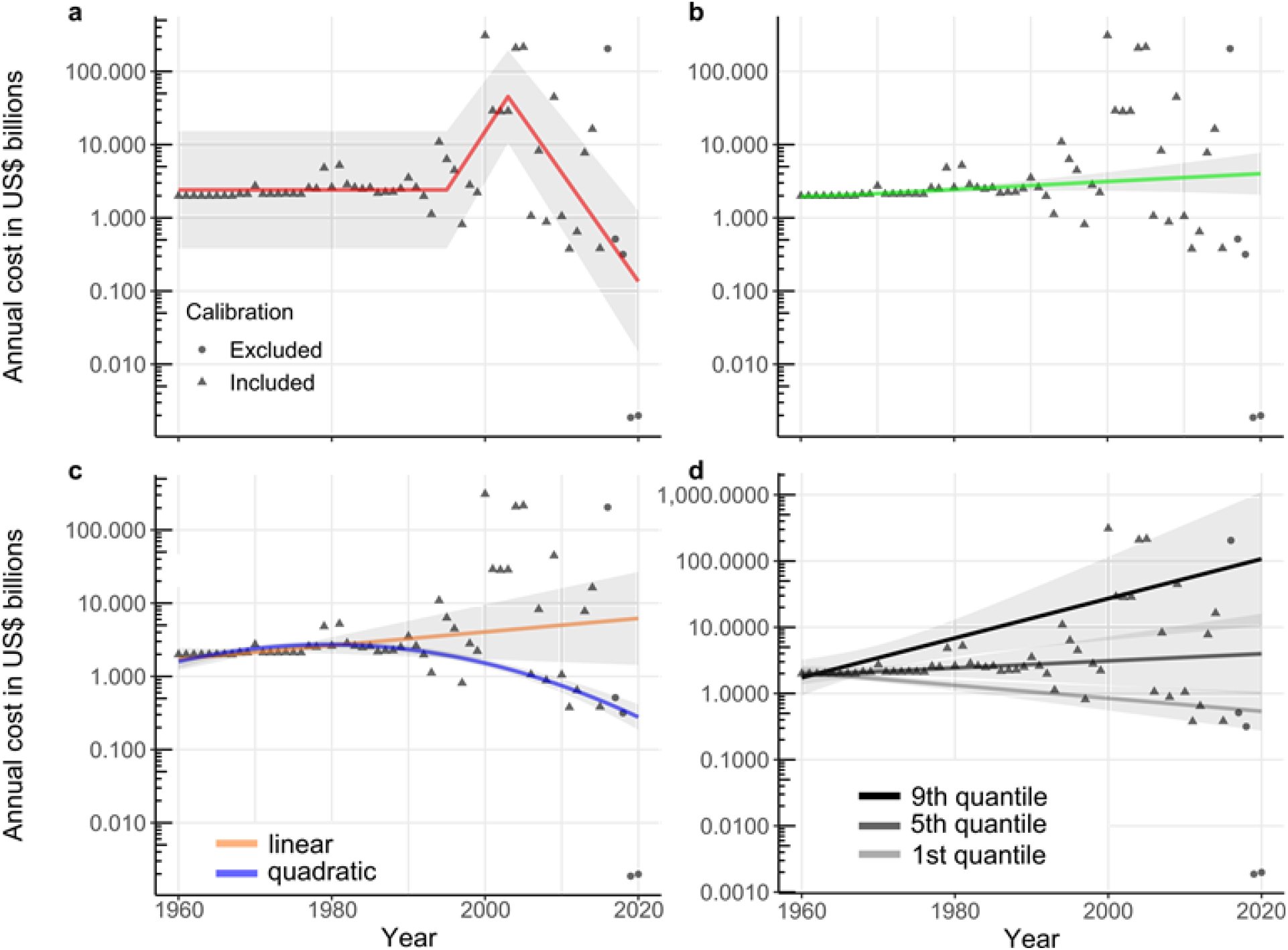
Statistical trends over time (1960–2016) of highly reliable, observed annual invasion costs recorded in the United States.The last four years of costs are excluded from the analysis due to time lags in data reporting causing <50% of costs to be reported. In spite of removing the last four years, substantial numbers of costs are likely still unreported, and apparent decreases in the non-linear models are driven by incomplete costs in recent years depressing the annual costs. (a) generalized additive model (GAM; green), (b) robust regression (linear: orange; quadratic: blue), and (c) quantile regressions. Error bands represent 95% confidence around the estimate.

## Discussion

Biological invasions have cumulatively caused at least $1.22 trillion in observed, highly reliable economic losses in the United States over the past six decades, with the largest impacts coming from mammalian and insect invaders. Agriculture suffered the highest costs, reflecting this sector’s high susceptibility to economic damage from non-native species (Paini et al. 2016). The predominance of terrestrial systems in reported economic costs is surprising given the importance of aquatic systems for ecosystem services and livelihoods (Darwall et al. 2018), but may follow the large damages to agriculture and could reflect the wider focus within ecology towards terrestrial ecosystems (Menge et al. 2009) and a relative lack of economic assets in aquatic realms. The fact that over two-thirds of observed costs were damages and losses is significant given these costs are harder to observe at large scales and more likely than management costs to be classified as potential (and thereby excluded from this analysis in the interest of conservatism). This indicates there may be considerable gains to be made from increased spending on biosecurity and post-invasion management. For example, the US Department of Interior reported spending only $117 million to manage invasive species (United States Department of the Interior 2020), despite managing approximately 21% of the area of the United States (United States Department of the Interior 2019). Of the money spent on management, only $1.35 billion (1% of $119.05 billion) is spent on pre-invasion biosecurity, despite work highlighting the cost-effectiveness of biosecurity protocols over longer-term management strategies (Leung et al. 2002, Lodge et al. 2006; Ahmed et al. this issue). Future investments in preventative measures and surveillance could help to offset future control and eradication costs in the United States and make those expenditures more effective in reducing damage and loss.

Costs from invasions appear to have unequal regional distribution, with the West reporting the greatest region-specific economic impacts and the Northeast the least. It is unclear whether the region-specific impacts in the West represent a distinct set of damaging species, better cost reporting, or a combination of the two. We observe that individual regions exhibited distinct patterning in costliest taxa and impacted sectors, potentially representing disparate cost reporting at the national scale, or differences in invasion patterns, introduction pathways and economic or environmental contexts. Notably, the costliest species US-wide did not appear as the costliest species within regions, indicating the species with greatest impact nationwide may not be the same as those within a region. While the region-specific costs provide important insight into how risks and costs differ among regions, most economic costs lacked the spatial resolution necessary to be attributed to a specific US state or region. We think that efforts to improve resolution and standardization of cost reporting would provide a clearer picture of overall costs and allow for more efficient mobilization of funding at relevant spatial scales.

Nationally and regionally, reported invasion costs were dominated by animals. Terrestrial animals, particularly mammals and insects, include some of the most notorious invasive species, and the large reported costs are a combination of substantial damages, management costs, and study effort associated with these species. For example, the two mammal species with the largest reported costs are feral cats and black rats, which inspire extensive research effort due to their ecological impacts (Loss et al. 2012, Knowlton et al. 2007), often close proximity with human populations, and the fact that both species inspire strong feelings, if for different reasons (Bjerke and Østdahl 2004, Hall et al. 2016, Jaric et al. 2020). At the same time, insects caused considerable damage to US forestry, agriculture, and health sectors, as has been shown to be the case globally (Bradshaw et al. 2016). Climate change may exacerbate such costs; in particular, it has been shown that it could lead to an average increase of 18% in areas suitable for global arthropod invaders (Bellard et al. 2013). While our findings match some other studies (e.g., Pimentel et al. 2000, 2005, Bradshaw et al. 2016) in terms of costliest taxa, this may not hold true in terms of ecological impact. In fact, the three US invasive species most frequently studied in terms of ecological impact were red swamp crayfish (*Procambarus clarkii*), the red imported fire ant, and Amur Honeysuckle (*Lonicera maackii*; Crystal-Ornelas et al. 2020). However, robust estimates of economic costs are only available for red imported fire ant (e.g., Lard et al. 2001). Further, due to a lack of published reports of their economic impacts, many well-known aquatic invaders such as the six problematic Asian carp species (Family *Cyprinidae*), northern snakehead (*Channa argus*), and rusty crayfish (*Orconectes rusticus*) had few, if any, cost estimates in our database. Though it is reasonable to assume that well-studied species also have significant economic impact, it should be noted that even poorly studied species may have similar effects and the majority of invasive species lack economic cost appraisals at national scales (e.g., Gren et al. 2009).

Our estimate of annual costs due to invasive species in the United States is lower than the $120-138 billion annually often cited from earlier studies (Pimentel et al. 2000, 2005), but our approach and methods make this estimate more comprehensive and detailed. For example, our focus on highly reliable, observed costs enables costs to be assigned to specific time periods, in contrast to unspecified time frames in the Pimentel papers (Pimentel et al. 2000, 2005). As a result, we have used the best available data to determine a lower floor for the reported costs of invasive species in the United States. When we include potential and lower reliability cost entries in our calculations, the total cost increases to $4.52 trillion, or $ 74.10 billion annually, while still relying on costs assigned to defined time periods. Additionally, there are several reasons why our calculated costs are likely substantial underestimates, particularly in recent years, where extremely high variance limits confidence in the estimate of an annual average cost. Much of this variance results from two countervailing forces in the data. First, when reports failed to define the actual time period over which costs result, we conservatively assigned these costs to a single year (see Methods, Diagne et al. 2020a), potentially inflating costs in some years and likely underestimating them in other years. Related complications in accurately calculating costs arise from the difficulty of extrapolating costs over the full period of species impacts. Second, pervasive time lags between cost occurrences and their reporting reduced costs in recent years, and we had limited ability to accurately correct for this, other than as we did, by omitting the most recent, and probably amongst the costliest, four years. Additionally, if economic impacts of invasions follow time lag patterns similar to ecological impacts (Essl et al. 2011), the true economic costs of current invasions may take decades to manifest.

Additional missing costs result from the inherent difficulty in estimating economic losses in relation to ecosystem services (Nunes et al. 2001; but see Hanley 2019). While these non-market-type costs are not captured in our current analysis if not monetarily valued, the $102.46 billion in damage/losses found in the environmental sector may serve as an indicator of these sorts of losses. Underestimation also stems from gaps in accounting of both the known and unknown damage caused by some taxonomic groups and impacted habitat types. Furthermore, non-market costs (e.g., nutrient cycling, water filtration, etc.) are rarely assigned value and can be substantially higher than market costs (Holmes et al. 2009). Finally, economic impacts for most invasive species have simply not been estimated (Gren et al. 2009, Cuthbert et al. 2021), which may reflect broader biases in ecological impact research across habitats and geographic regions (Crystal-Ornelas 2020). Such biases may explain why costs of terrestrial invaders were greater than aquatic invaders, despite the often-cited impacts of the latter (Ricciardi et al. 2011). Given the missing and biased costs, the figures presented here within should not be considered as static, final amounts but rather as the most current and inclusive estimate of the minimum costs of invasions in the United States thus far. Indeed, these outcomes are the results of analyses of a database that is expected and intended to evolve over time, and which will offer unique opportunities to improve and refine cost information (Diagne et al. 2020b).

In the past six decades, US invasion costs have apparently increased as a result of both increasing invasion rates and spread of extant invasions. Although a couple of our models exhibit declining trends in invasive costs, these models appear to be spuriously leveraged by missing data in the most recent and, likely, costliest years. Thus, we believe that the models illustrating increasing costs over time are the most probable. In that regard, the GAM and linear robust regression, which were less sensitive to outliers, predicted reported invasion costs in 2020 to be at least $4.01 and $6.19 billion, respectively. Indeed, across the globe, invasion rates show no signs of saturation (Seebens et al. 2017). In the future, as global trade increases and climate change patterns continue to intensify (Seebens et al. 2018, Bellard et al. 2013), we anticipate a further increase in invasion costs, particularly if investments in biosecurity remain insufficient to prevent future introductions (Leung et al. 2002). Given these increasing numbers and the lack of cost information we have for most invaders, our findings provide critical information for managers, planners, and policymakers. We urge increased and improved (i.e., standardized, Diagne et al. 2020a) cost reporting by stakeholders and managers in the context of biological invasions. An integrated data collection point, such as the InvaCost database, is an important first step in offering future decision-makers a comprehensive approach to report and utilize estimates of economic costs. Future works should seek to rectify existing knowledge gaps in economic costs across taxonomic, geographic, and environmental scales, in the United States and beyond. Furthermore, increased investments in biosecurity to reduce arrival and secondary spread of non-natives is urgently needed if costs are to be contained. Given that proactive management can be magnitudes more cost-effective than ongoing damages (Leung et al. 2002; Ahmed et al., this issue), monitoring, biosecurity and control investments should be prioritized to offset future costs and societal disruptions from invasions. Ultimately, our work provides an urgent warning on the massive, expanding costs of invasive species across the US that if left unchecked will continue unabated.

## Acknowledgments

The authors wish to thank all InvaCost workshop attendees and project participants who contributed to the development of the InvaCost database. PJH would like to thank his daughters who kept him awake at night and expresses his gratitude to the existence of energy drinks which got him through the analysis of the data.

## Declarations

### Data and materials availability

All data needed to evaluate the conclusions in the paper are present in the paper and/or the Supplementary Materials.

### Code availability

Links to all code needed to evaluate conclusions are present in the paper

### Author contributions

This work is the result of a workshop in which all authors actively participated. FC and CD designed the research, and JFL coordinated it; CD and FC developed and managed the database and JFL, PJH, AMK and RNC updated and cleaned the US subset of the database; JFL, PJH, AMK, RNC and AJT analyzed the data; JFL, PJH, AMK and RNC wrote the first draft of the paper, with further input from all authors. All authors read and approved the final manuscript.

## Supplementary Materials

**Fig. S1.**
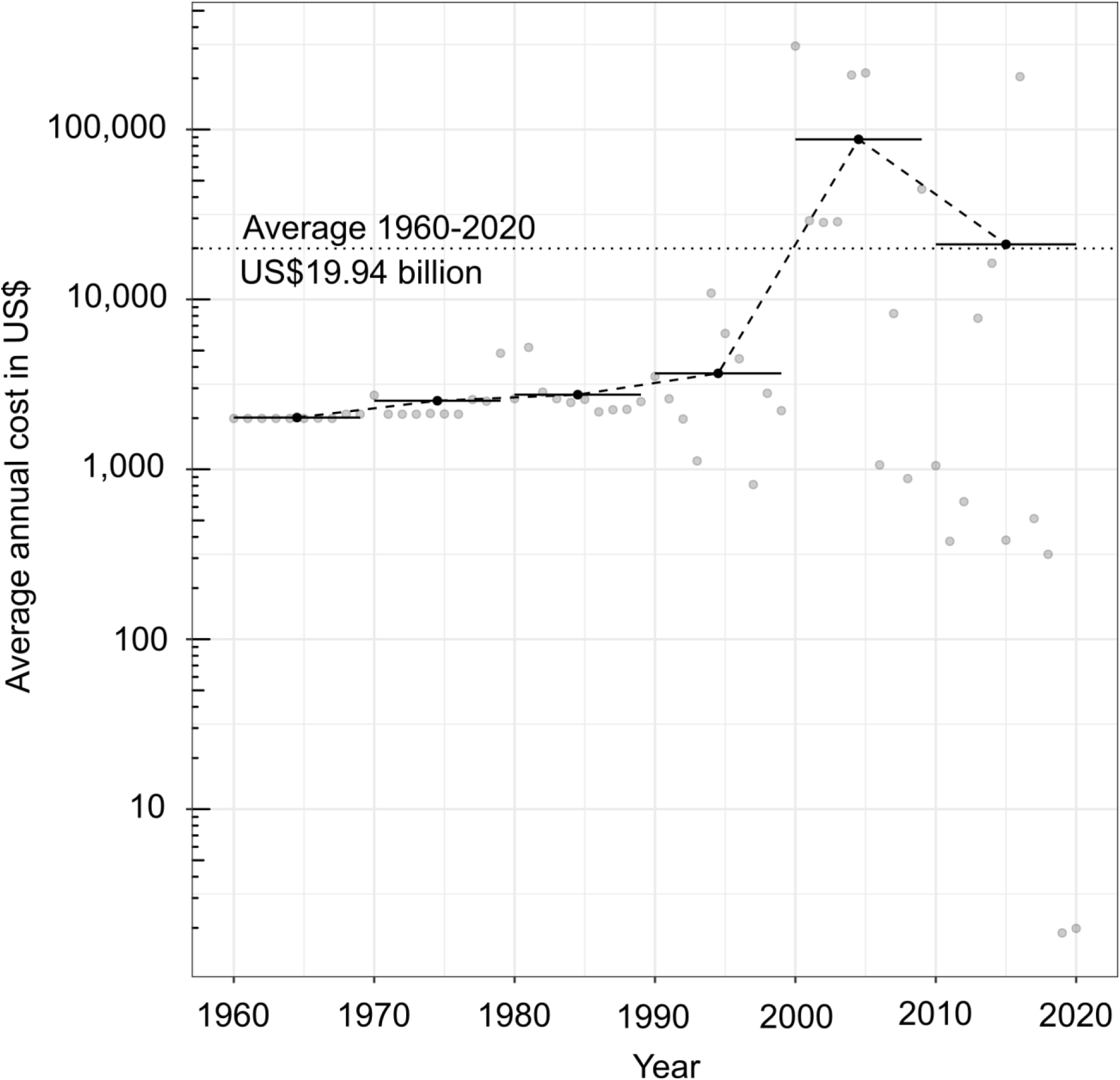
Annual cost accumulations of invasive species in the United States between 1960 and 2020.

**Table S1.** Distributions of cost totals and database entries across regions and sectors. Totals are from the expanded database.

| Region | Impacted sector | Cost (\$ billion) | Database entries |
| --- | --- | --- | --- |
| Northeast | Agriculture | 0.003 | 3 |
| Southeast | Agriculture | 0.488 | 17 |
| Midwest | Agriculture | 0.695 | 14 |
| West | Agriculture | 16.567 | 70 |
| Southwest | Agriculture | 0.751 | 11 |
| Outlying territories | Agriculture | 0.002 | 5 |
| Multiregional/unspecified | Agriculture | 491.047 | 139 |
| Northeast | Authorities/stakeholders | 0.119 | 59 |
| Southeast | Authorities/stakeholders | 0.870 | 110 |
| Midwest | Authorities/stakeholders | 0.079 | 17 |
| West | Authorities/stakeholders | 6.656 | 376 |
| Outlying territories | Authorities/stakeholders | 0.032 | 45 |
| Multiregional/unspecified | Authorities/stakeholders | 29.350 | 232 |
| Southeast | Environment | 0.016 | 8 |
| Midwest | Environment | 0.003 | 3 |
| Southwest | Environment | 0.003 | 3 |
| Multiregional/unspecified | Environment | 102.431 | 52 |
| Southeast | Fisheries | 0.031 | 3 |
| Midwest | Fisheries | 0.002 | 1 |
| Multiregional/unspecified | Fisheries | 0.007 | 3 |
| Northeast | Forestry | 0.194 | 1 |
| Southeast | Forestry | 15.532 | 2 |
| Multi/unspecified | Forestry | 26.872 | 15 |
| Southeast | Health | 0.008 | 54 |
| Outlying territories | Health | 0.000 | 2 |
| Multiregional/unspecified | Health | 19.481 | 5 |
| Northeast | Public/social welfare | 0.312 | 2 |
| Southeast | Public/social welfare | 0.064 | 6 |
| Midwest | Public/social welfare | 0.024 | 14 |
| West | Public/social welfare | 1.647 | 100 |
| Southwest | Public/social welfare | 1.304 | 12 |
| Outlying territories | Public/social welfare | 0.190 | 11 |
| Multiregional/unspecified | Public/social welfare | 37.194 | 21 |
| Northeast | Multisectoral/unspecified | 0.005 | 12 |
| Southeast | Multisectoral/unspecified | 0.295 | 11 |
| Midwest | Multisectoral/unspecified | 0.304 | 10 |
| West | Multisectoral/unspecified | 3.753 | 149 |
| Outlying territories | Multisectoral/unspecified | 0.816 | 31 |
| Multiregional/unspecified | Multisectoral/unspecified | 458.918 | 69 |

**Table S2.**
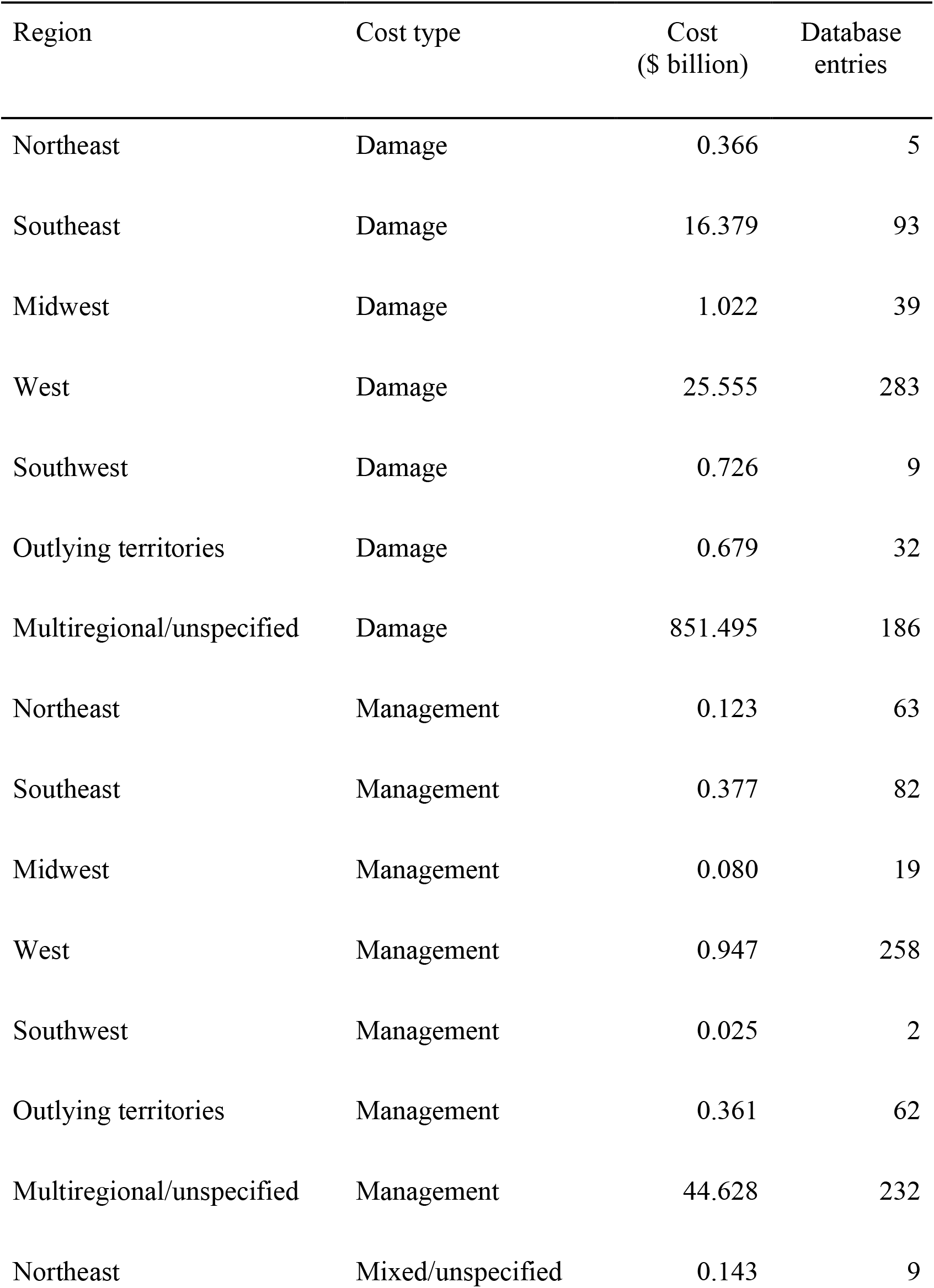
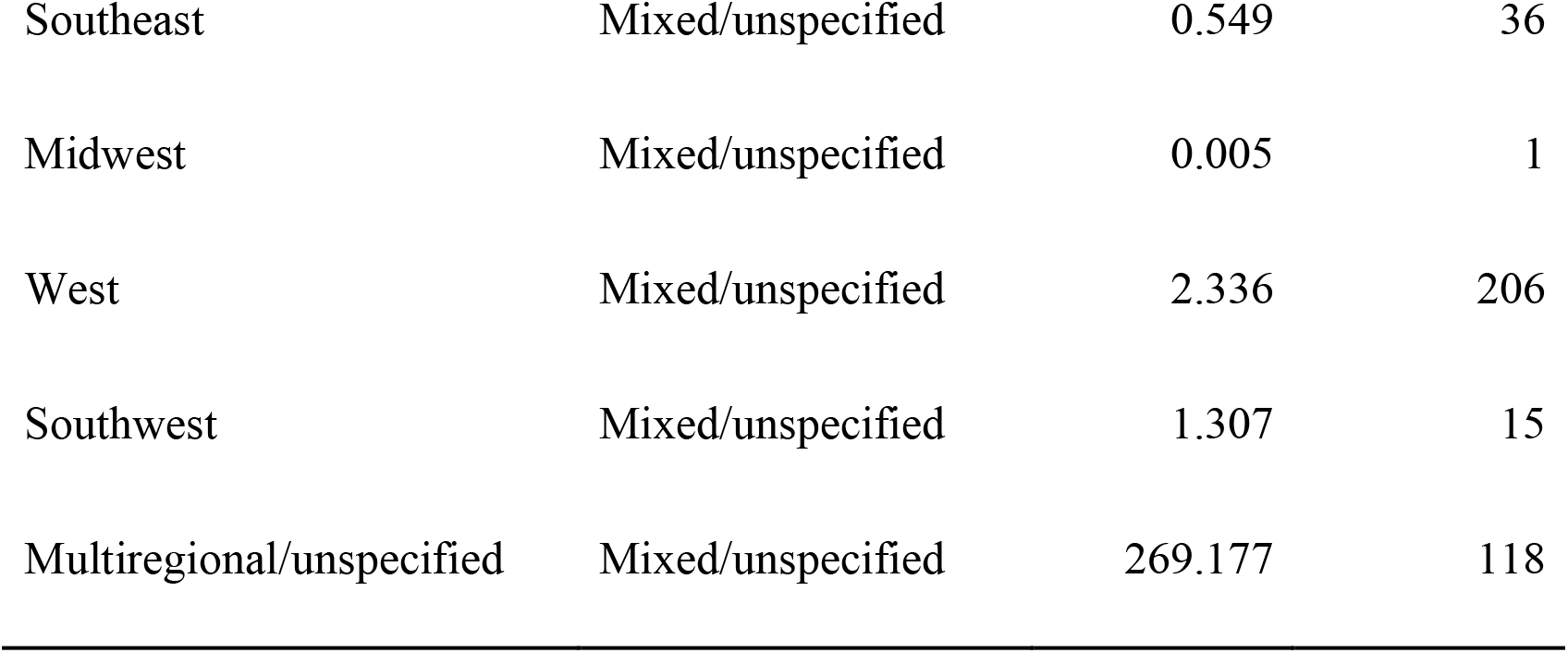
Distributions of cost totals and database entries across regions and cost types. Totals are from the expanded database.

| Region | Cost type | Cost<br>(\$ billion) | Database<br>entries |
| --- | --- | --- | --- |
| Northeast | Damage | 0.366 | 5 |
| Southeast | Damage | 16.379 | 93 |
| Midwest | Damage | 1.022 | 39 |
| West | Damage | 25.555 | 283 |
| Southwest | Damage | 0.726 | 9 |
| Outlying territories | Damage | 0.679 | 32 |
| Multiregional/unspecified | Damage | 851.495 | 186 |
| Northeast | Management | 0.123 | 63 |
| Southeast | Management | 0.377 | 82 |
| Midwest | Management | 0.080 | 19 |
| West | Management | 0.947 | 258 |
| Southwest | Management | 0.025 | 2 |
| Outlying territories | Management | 0.361 | 62 |
| Multiregional/unspecified | Management | 44.628 | 232 |
| Northeast | Mixed/unspecified | 0.143 | 9 |
| Southeast | Mixed/unspecified | 0.549 | 36 |
| Midwest | Mixed/unspecified | 0.005 | 1 |
| West | Mixed/unspecified | 2.336 | 206 |
| Southwest | Mixed/unspecified | 1.307 | 15 |
| Multiregional/unspecified | Mixed/unspecified | 269.177 | 118 |

**Table S3.** Distributions of cost totals and database entries across regions and environments. Totals are from the expanded database.

| Region | Environment | Cost<br>(\$ billion) | Database<br>entries |
| --- | --- | --- | --- |
| Northeast | Terrestrial | 0.299 | 34 |
| Southeast | Terrestrial | 16.774 | 68 |
| Midwest | Terrestrial | 1.097 | 46 |
| West | Terrestrial | 21.924 | 525 |
| Southwest | Terrestrial | 2.058 | 26 |
| Outlying territories | Terrestrial | 0.038 | 55 |
| Multiregional/unspecified | Terrestrial | 601.322 | 402 |
| Northeast | Semi-aquatic | 0.004 | 4 |
| Southeast | Semi-aquatic | 0.130 | 89 |
| West | Semi-aquatic | 0.198 | 44 |
| Outlying territories | Semi-aquatic | 1.002 | 39 |
| Multiregional/unspecified | Semi-aquatic | 0.235 | 17 |
| Northeast | Aquatic | 0.324 | 37 |
| Southeast | Aquatic | 0.364 | 45 |
| Midwest | Aquatic | 0.005 | 12 |
| West | Aquatic | 1.603 | 69 |
| Multiregional/unspecified | Aquatic | 11.151 | 26 |
| Northeast | Mixed/unspecified | 0.004 | 2 |
| Southeast | Mixed/unspecified | 0.037 | 9 |
| Midwest | Mixed/unspecified | 0.005 | 1 |
| West | Mixed/unspecified | 5.112 | 109 |
| Multiregional/unspecified | Mixed/unspecified | 552.592 | 91 |

**Table S4.** Species with highest reported costs in the United States. Geographic origin, and number of database entries is included.

| **Rank** | **Species** | **Geographic origin** | **Cost**<br>**(\$ billion)** | **Database**<br>**entries** |
| --- | --- | --- | --- | --- |
| 1 | *Felis catus* | Africa (Egypt) | 45.54 | 6 |
| 2 | *Rattus rattus* | Asia (India) | 23.84 | 3 |
| 3 | *Coptotermes formosanus* | Asia (China) | 18.53 | 7 |
| 4 | *Anthonomus grandis* | Mexico | 9.67 | 98 |
| 5 | *Wasmannia auropunctata* | Central and South America | 7.31 | 27 |
| 6 | *Apis mellifera* | Asia | 5.68 | 3 |
| 7 | *Solenopsis invicta* | South America | 5.01 | 46 |
| 8 | *Ophiostoma ulmi* | Asia | 3.64 | 9 |
| 9 | *Rhipicephalus microplus* | Asia | 3.55 | 1 |
| 10 | *Lymantria dispar* | Eurasia and Asia | 3.54 | 76 |

**Table S5.** Root mean square errors (RMSE) across models. These consider both adjusted (accounting for time lags) and full data. Abbreviations: OLS = ordinary least squares, RR = robust regression, MARS = multiple additive regression splines, GAM = generalized additive model, QT = quantile

|  | RMSE<br>(calibration) | RMSE<br>(all data) |
| --- | --- | --- |
| OLS (linear) | 0.58 | 0.90 |
| OLS (quadratic) | 0.57 | 0.87 |
| RR (linear) | 0.58 | 0.89 |
| RR (quadratic) | 0.73 | 0.87 |
| MARS | 0.47 | 0.66 |
| GAM | 0.59 | 0.87 |
| QT (0.1) | 0.83 | 0.97 |
| QT (0.5) | 0.59 | 0.87 |
| QT (0.9) | 0.83 | 1.25 |

**Dataset S1 (separate file). Subset of the InvaCost database for the United States and its territories.** The database contains 1404 entries specific to the United States, derived from 407 studies.

